# Quantified duplications of proteins within complexes across eukaryotes

**DOI:** 10.64898/2026.02.23.705090

**Authors:** Ore Francis

**Affiliations:** Investigation of protein complex orthology

## Abstract

Protein complexes are central to cell biology and typically verified via a combination of interaction data, complete genome sequencing and comprehensive protein-coding gene predictions for reference eukaryotes. However this data is lacking for non-reference eukaryotes.

Protein complexes can be predicted in species for which no interaction data is available by mapping orthology of verified protein complex components from reference eukaryotes to predicted proteomes. Studies that map conservation of protein complex components by orthology are often limited to a small number of protein queries, an under-representation of non-reference, microbial eukaryotes and are scattered across the literature. Here, I integrate orthology and protein interaction data by mapping proteins of experimentally verified complexes to orthogroups of proteins spanning 31 diverse eukaryotes. Proteins within complex-harbouring orthogroups are retained and distributed more evenly across taxa than non-complex orthogroups. I identified 184 universal orthogroups that included orthologs of known protein complex components from all 31 eukaryotes, consistent with a conserved core repertoire, likely present in the last eukaryotic common ancestor (LECA).

I generated the protein complex orthology cartographer (PCOC) suite to find significant duplications and reductions of proteins in universal orthogroups across and between eukaryotes. This revealed both multi-copy and notably single-copy proteins, in all queried species, from the exosome, spliceosome, proteasome, small-ribosomal processome, tRNA synthetases, MCM complexes and RNA polymerase III. Case analyses of Naegleria gruberi and Guillardia theta highlight taxon-specific expansions and show how broader protist inclusion improves domain-wide inference of eukaryotic protein-complex evolution.

## Introduction

Proteins underpin the majority of cellular functions across prokaryotes, eukaryotes and viruses. While proteins regularly function together with nucleic acids, carbohydrates and lipids, they also interact with other proteins in protein complexes. For example, at least 30% of the known human proteome form stable complexes (Steinkamp *et al*., 2025). The formation of a protein complex typically refers to a specific, non-transient physical association of two or more proteins, facilitated by the physical properties of protein domains to form stable and specific interactions with others (Spirin and Mirny, 2003; Pereira-Leal *et al*., 2007). Protein complexes are ubiquitous components of all cellular life, from components of extant organelles to those present in the last eukaryotic common ancestor. Accordingly, proteins that form protein complexes of organelles or are involved in cellular processes that are conserved throughout eukaryotes can serve as useful markers for comparative-genomic reconstruction of ancestral forms (Koonin, 2010; Sebé-Pedrós *et al*., 2014; More *et al*., 2020; More *et al*., 2024). Importantly, only a small number of proteins are expected to originate from horizontal gene transfers and therefore should not be included in ancestral reconstructions (Van Etten and Bhattacharya, 2020).

To determine unambiguous presence or absence of protein complexes across the proteome of a species, the genome of the species must first be sequenced. Furthermore, deep comparative-genomic reconstruction of eukaryotic protein complexes requires broad taxonomic representation across the domain, that can only be achieved by selecting taxa from the largest, monophyletic, and often exclusively protist taxonomic groups, known as eukaryotic supergroups (Burki *et al*., 2007; Koonin, 2010). Therefore, one challenge complicating comparative analysis of eukaryotic protein complexes is that proteome-wide protein complex prediction is only possible for 24,000 species (Challis *et al*., 2023), from the 2 million named (Blaxter *et al*., 2022) and potentially hundreds of millions more presumed (Mora *et al*., 2011; Wiens, 2023) extant eukaryotes. A further challenge is that only 1301 protists have been sequenced, from over 70,000 named and up to 10 million presumed extant protists (Adl *et al*., 2007; Challis *et al*., 2023).

For in silico comparative analyses of protein complexes, individual complexes (Francis, Han and Adams, 2013) or complexes involved in a particular process are typically studied (More *et al*., 2020), often including careful selection of the few available protist sequences that span known eukaryotic genome diversity. Multiple studies of protein complex orthology that primarily use protist sequences have demonstrated domain-wide evolution of macromolecular protein complexes across eukaryotes. Wideman recovered previously unknown patterns of conservation of endoplasmic reticulum (ER) membrane protein complex (EMC) components in 45 protist lineages (Wideman, 2015). More and colleagues collated a number of examples of membrane trafficking proteins, many of them complex components, homologs of which have been identified in protist lineages (More *et al*., 2020).

Ongoing improvements in sampling and sequencing technologies are producing an increasing number of eukaryotic genomes, gene and protein sequences. Automated annotation of sequence data with protein interaction results are gradually increasing the availability of protein-protein interaction data necessary to predict the protein complexes of eukaryotes. For example, a comprehensive database of protein interactions can be accessed via the IntAct portal (Toro *et al*., 2022), documenting more than 1.75 million binary interactions between proteins, combining physically determined interactions and those inferred from colocalisation. Tools and formats, such as fluorescence-resonance energy transfer (FRET) microscopy, comigration assays or co-association in microscopy are used to demonstrate protein-protein interactions but are not usually sufficient to infer participation of interacting proteins in complexes. Most physical associations of protein complexes are identified through immunoprecipitation, pull-down, tandem affinity purification, yeast-2-hybrid and cross-linking experiments, typically using a small number of query proteins.

The EBI Complex Portal, a related repository of protein complex data, relies on the expertise of curators to manually validate predicted protein complexes using a mixture of resources that include physical interaction and orthology data (Meldal *et al*., 2019). The extent of conservation of protein complexes across eukaryotes is likely significant. The most abundant, functional Gene Ontology (GO) term annotations for EBI Complex Portal proteins are “catalytic activity (28.1%)” and “binding (66.7%)” corresponding to broad enzymatic classification and protein-protein interaction ability, respectively. Broadly, 20% of proteins in the entire Universal Protein Knowledge Base (UniProtKB) that have GO annotations also are annotated as “binding” which suggests that existing data can assign over 52 million proteins to protein complexes (Apweiler *et al*., 2004; The Gene Ontology Consortium *et al*., 2023).

A wealth of sequencing data, gene predictions and orthology mappings are available for reference organisms. However complete protein complex repertoires - also termed complexomes - are rare. Few studies profile the entire repertoire of protein complexes for a eukaryotic cell, particularly in microbial eukaryotes. The concept of a protein complexome is relatively recent, with the early collation of protein complexes from *Saccharomyces cerevisiae* serving as the first notable example, pioneering the availability of a yeast complexome (Deshaies *et al*., 2002; Meldal *et al*., 2021). De novo complexomic experiments are rare, expensive and restricted to taxonomically narrow reference organisms. Therefore, only a small number of complexomes have been published and added to molecular interaction databases since 2002 and attempts to unify this data are very rare (Havugimana *et al*., 2012; Wan *et al*., 2015; Drew *et al*., 2017; McWhite *et al*., 2020; Skinnider *et al*., 2021).

One approach to unifying such data was the compilation of an ancestral eukaryotic protein complexome, inferred from genomes of animals, plants and fungi (Ceulemans, Beke and Bollen, 2006). Ceulemans et al compiled a list of experimentally verified protein complexes from the literature and determined homologs in animal, plant and fungi genomes. However, corresponding protist sequences were not available at the time. Nevertheless they defined an ancestral protein complex repertoire, using the concept of a Eukaryotic Virtual Ancestor (EVA). EVA comprised 1371 sequences and they noted remarkable complex component loss in the microsporidian *Encephalitozoon cuniculi*. As well as manually resolving some paralogy issues, Ceulemans and colleagues grouped co-orthologs together (Ceulemans, Beke and Bollen, 2006). Today, standardised orthology solutions such as the Orthologous (OrthoDB) and Orthologous Matrix (OMADB) databases are viable solutions to mapping genome-wide protein predictions across the domain of eukaryotes (Simão *et al*., 2015; Altenhoff *et al*., 2024; Karasikov *et al*., 2024).

In this study I investigate conservation of protein complexes across eukaryotes by mapping known eukaryotic protein complexes to the genomes of 31 eukaryotes that span 11 supergroups. I show that orthogroups containing complex-forming proteins are more evenly distributed across taxa than the other orthogroups, suggesting underlying pressure to conserve protein complexes.

By inferring how many orthologs of complex-forming proteins are encoded in each genome of 31 species that span eukaryote diversity, I determine which orthologs are likely conserved in LECA, where taxon-specific expansions and reductions of these orthologs occur across the queried genomes and which known heterodimeric-interaction do these expansions and reductions correspond to. I highlight expansions and reductions identified in two protists, *Naegleria gruberi*, an excavated protist, similar to LECA, and the four genome-harbouring *Guillardia theta* (FritzLaylin *et al*., 2010; Curtis *et al*., 2012). By defragmenting protein interaction and orthology data, this method may be valuable as a metric in the assessment of future novel, divergent eukaryote genomes.

## Methods

### UniProt searches

A representative set of eukaryote genomes corresponding to the animal, plant, fungi and protist species was selected to determine conservation of protein complexes across eukaryotes. Taxa were selected to span the diversity of eukaryotes, including a majority of protists. To develop a framework based on OrthoDB orthology data (Tegenfeldt *et al*., 2025), taxa included in OrthoDB version 12 (odb12v0) were selected. The total number of genomes selected was 31, achieving representation of taxa across opisthokonta (Metazoa, Fungi and Choanoflagellates), Amoebozoa, Stramenopiles, Alveolates and Rhizaria (the SAR group), Viridiplantae (Chlorophyta and Streptophyta), Haptophyta, Cryptophyta and Heterolobsea supergroups, without over-representation of any individual group. Full taxonomic subset is available in Tab. 1.

**Table 1:** Taxonomic subset used in this study.

| Taxonomy ID | Species name |
| --- | --- |
| 484906 | *Babesia bovis* |
| 333668 | *Theileria parva* |
| 508771 | *Toxoplasma gondii* |
| 5888 | *Paramecium tetraurelia* |
| 312017 | *Tetrahymena thermophila* |
| 403677 | *Phytophthora infestans* |
| 2880 | *Ectocarpus siliculosus* |
| 37360 | *Plasmodiophora brassicae* |
| 296587 | *Micromonas commoda* |
| 3055 | *Chlamydomonas reinhardtii* |
| 3218 | *Physcomitrium patens* |
| 3702 | *Arabidopsis thaliana* |
| 3988 | *Ricinus communis* |
| 905079 | *Guillardia theta* |
| 280463 | *Emiliana huxleyi* |
| 284813 | *Encephalitozoon cuniculi* |
| 645134 | *Spizellomyces punctatus* |
| 578462 | *Allomyces macrogynus* |
| 4932 | *Saccharomyces cerevisiae* |
| 330879 | *Aspergillus fumigatus* |
| 4896 | *Schizosaccharomyces pombe* |
| 237631 | The following |
| 9606 | *Homo sapiens* |
| 7227 | *Drosophila melanogaster* |
| 10228 | *Trichoplax adhaerens* |
| 6087 | *Hydra vulgaris* |
| 595528 | *Capsaspora owczarzaki* |
| 431895 | *Monosiga brevicollis* |
| 352472 | *Dictyostelium discoideum* |
| 13642 | *Heterostelium pallidum* |
| 5762 | *Naegleria gruberi* |

The UniProt database (release 2025_04) was searched using the Uniprot.ws package version 2.48.0 (Carlson, 2017) for all eukaryotic proteins in the EBI Complex Portal (Meldal *et al*., 2019) using the terms “organism_id:2759” and “keyword:ComplexPortal”, to retrieve all corresponding accessions, gene IDs, protein names, review statuses, Complex Portal IDs and OrthoDB accession numbers.

### OrthoDB searches

All orthogroup analysis was performed using OrthoDB version 12 (Tegenfeldt *et al*., 2025). The file containing mappings of eukaryote genes to orthogroups was downloaded from OrthoDB (odb12v0_OG2genes.tab from odb12v0). All genes corresponding to taxon IDs of the 31 species of interest were extracted from this file.

### Commandline scripts

To parse data from the odb12v0 and EBI Complex Portal data from UniProt, I developed a suite of six scripts, that I call The Protein Complex Orthology Cartographer (PCOC). PCOC is available as individual R scripts, with a BASH script to parse the odb12v0 database and a python script to retrieve National Center for Biotechnology Information (NCBI) taxonomy newick files for submitted taxa. Documentation is available at https://github.com/Oreoluwa85/PCOC. Briefly:

- generate_compOG_repertoire.R - is a script that generates a linear compOG repertoire, restricting search space to the repertoire of 184 compOGs found originally. The data outlined in this paper can be retrieved by running this script on a taxon list of the 31 eukaryote subset
- generate_compOG_totals.R - is a script that generates a histogram corresponding to the total number of orthogroups found stratified by taxa along an NCBI retrieved tree. The tree is rendered as a gg-object and can be styled manually.
- generate_compOG_pair_repertoire.R - is a script that generates a compOG pair repertoire, from shared orthogroups of a taxon list. The reference eukaryotic compOG pair repertoire outlined in this study can be retrieved by running this script on a taxon list of the 31 eukaryote subset
- generate_compOG_pair_score.R - is a script that generates a compOG pair repertoire, from the number of orthogroups of a single taxon, to be compared to a reference compOG pair repertoire of a taxon list including that taxon.
- generate_odb12_tabfile.sh - is a BASH script that generates a tab file of every orthogroup corresponding to every taxon passed to the script in a single taxon list using grep. The tabfile can take a long time to generate.
- Get_taxid_newick.py - is a python3 script that generates an NCBI taxonomy tree from a taxon list, that is automatically ran after generate_odb12_tabfile.sh

An rmarkdown document, that renders this paper, is available on Github and includes required packages and code required to generate the figures submitted within this paper.

### GoaT searches

Numbers of published eukaryote genomes in text acquired by searching genomes on a tree (GoaT-goat.genomehubs.org) with the following queries: - All eukaryotes: “tax_tree(!eukaryota) AND tax_rank(species, subspecies, strain”. - Protists: “tax_tree(!metazoa,!embryophyta,!fungi) AND tax_rank(species,subspecies,species group,species subgroup)”

## Results

### Identifying universally conserved, complex-protein-harbouring orthogroups

To determine pan-eukaryotic, complex-protein-harbouring orthogroups (compOGs), without taxonomic over-representation of any eukaryotic supergroup, I extracted orthogroups of 31 diverse eukaryotes, animals, plants, fungi and protists, from the OrthoDB (version odb12v0) database (Tegenfeldt *et al*., 2025). Orthogroups that contained eukaryotic proteins from a single superphylum, phylum or any taxonomic rank narrower than domain (i.e. eukaryotes) were excluded. All other orthogroups that contained eukaryotic proteins from the EBI Complex Portal or proteins from any of the 31 species were retrieved from odb12v0.

A total of 392,850 proteins were retrieved that were distributed across 82,213 orthogroups. A total of 8,129 eukaryotic proteins from the EBI Complex Portal were distributed across 4,123 compOGs. By matching orthogroups from the taxonomic subset and from the EBI Complex Portal, a pan-eukaryotic, repertoire of 3,347 compOGs was retrieved (Fig 1, Tab. 2). The remaining 78,866 orthogroups, referred to as non-compOGs, did not contain complex component homologs.

**Figure 1.**
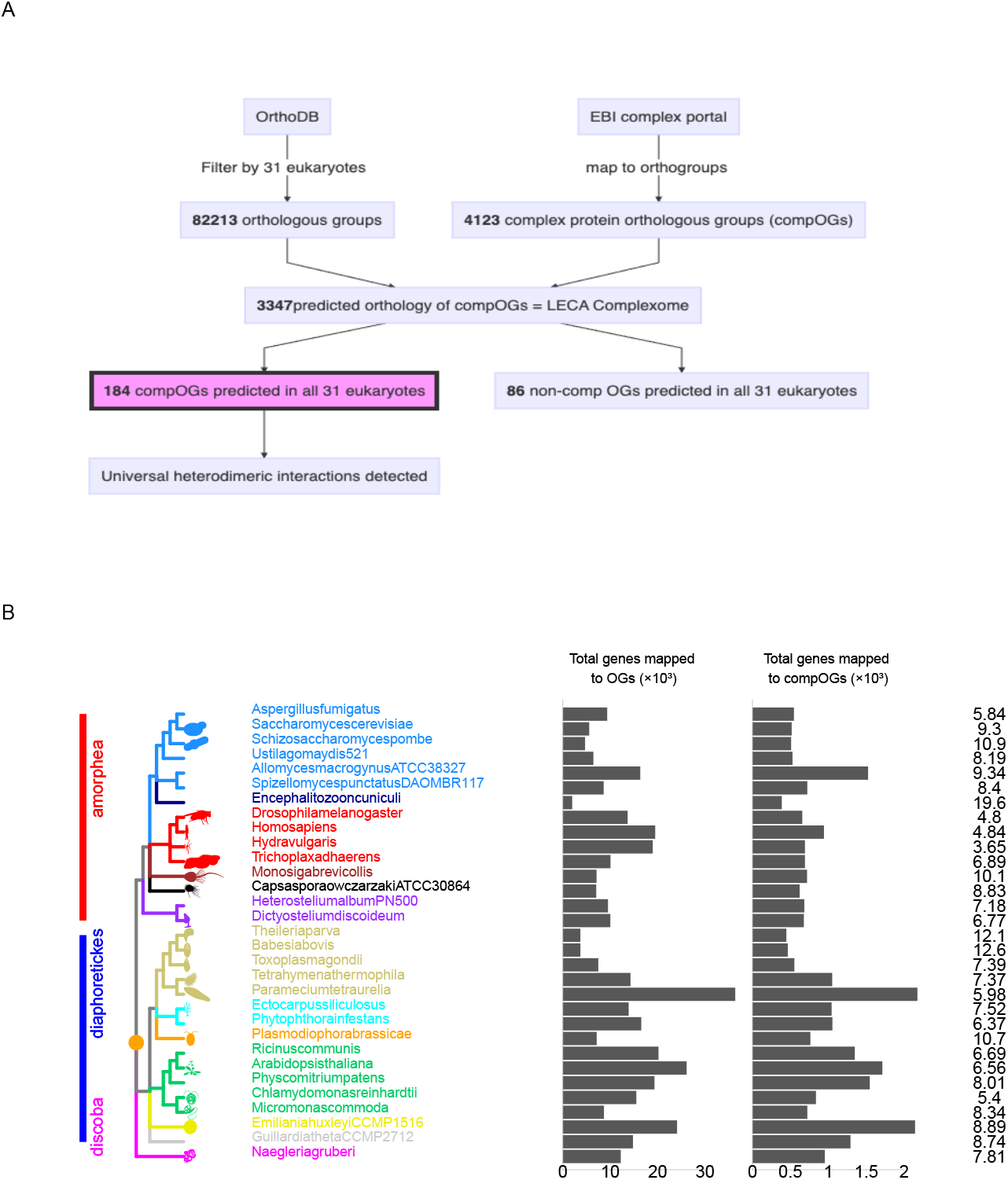
**A. Flowchart outlining filters implemented on database searches to isolate complex-protein-harbouring orthogroups (compOGs).** Numbers of orthogroups identified at each stage are in bold. **B. Summary of orthogroups identified in this study**. Numbers of total orthogroups and numbers of compOGs identified for each taxon shown as a bar graph. Percentage of total orthogroups that are compOGs for each taxon shown adjacent to each taxon-specific compOG bar.

**Table 2:** Numbers of proteins and corresponding orthogroups identified in Uniprot belonging to the taxa investigated in this study and in the EBI complex portal.

|  | Proteins | Orthogroups |
| --- | --- | --- |
| Euk subset | 392850 | 82213 |
| Complex Portal | 8929 | 4123 |
| Subset compOGs | 111844 | 3347 |
| Subset UcompOGs | 21377 | 184 |

*E. huxleyi, P. tetraurelia* and *A. thaliana* genomes mapped to more total orthogroups and more compOGs than other species. The expansion of proteins in some species may be driven by changes in ploidy: *P. tetraurelia* and *A. thaliana* have undergone multiple rounds of whole-genome duplication and *E. huxleyi* may have undergone ancestral genome duplications (Borkhardt and Olson, 1983; Simillion *et al*., 2002; Aury *et al*., 2006; Dassow *et al*., 2015; Carrier *et al*., 2018). *E. cuniculi* had the fewest number of orthogroups of any species in this study but the largest percentage of compOGs of any species (19.6%).

To compare how evenly compOG and non-compOG proteins are distributed across the subset of eukaryotes, both the coefficient of variation (the ratio of standard deviation to the mean) and the mean number of proteins per taxon were calculated for each compOG and non-compOG (Fig 2). Many highly-unevenly distributed orthogroups were retrieved, corresponding to, either species-specific proteins or proteins present in fewer than three species, despite restricting the taxonomic rank of all retrieved orthologous groups to domain level.

**Figure 2.**
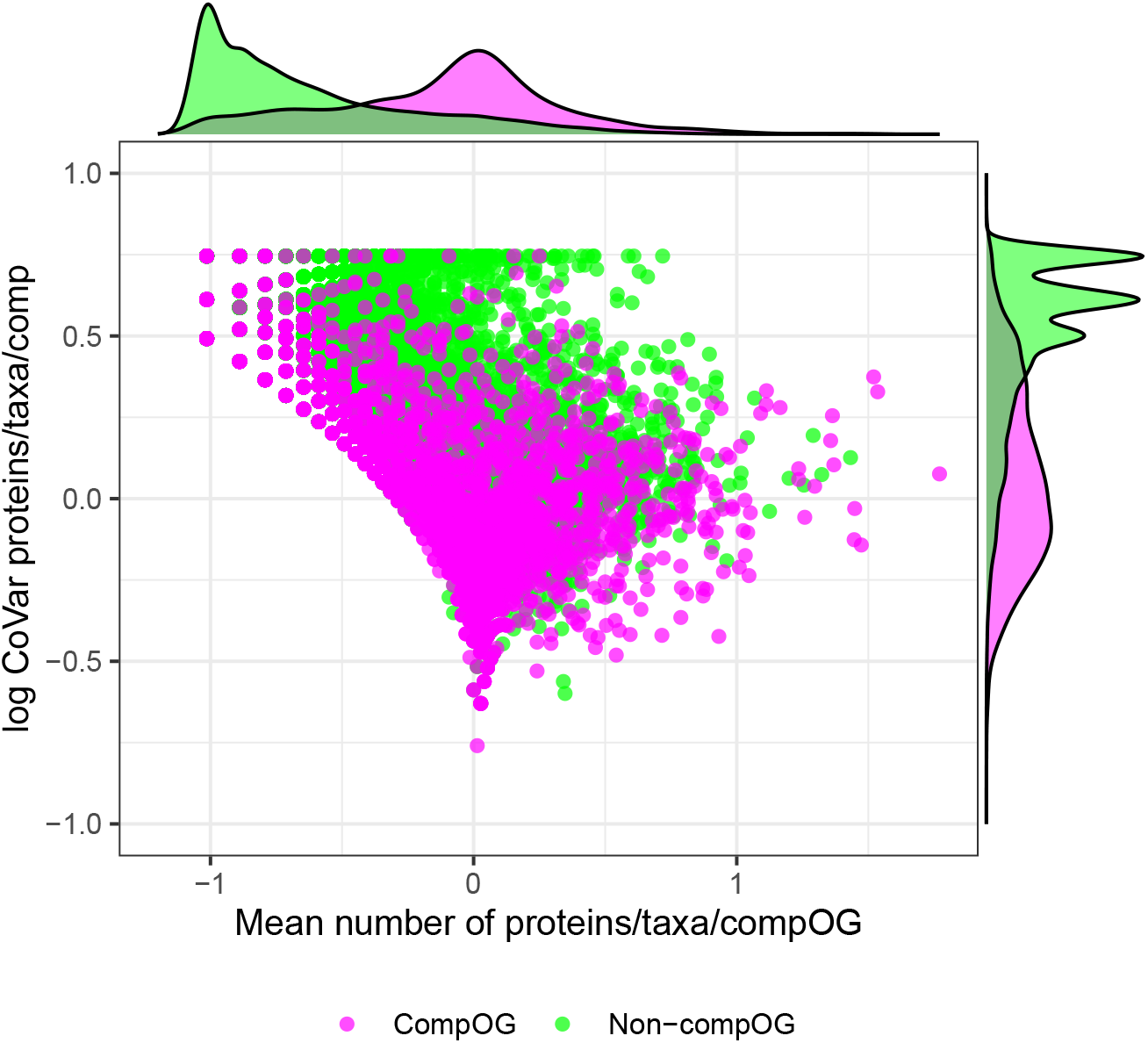
**A. Distribution of means for each orthogroup across 31 eukaryotes up to 0.1 (log(-1)).** Coefficient of variation of proteins per orthogroup between species for each orthogroup was plotted against mean value of numbers of proteins per orthogroup across species. Each green and magenta data point correspond to each non-compOG and compOG, respectively. Frequency distributions corresponding to log of the means and log of the coefficients of variation (CVs) of compOGs and non-compOGs are shown along the top and the right-hand sides of the plot, respectively. Frequency distributions show high density of non-compOGs with means well below one (non-proportional to B). **B Distribution of means for each orthogroup across 31 eukaryotes from 0.01 to 10000**. Frequency distributions demonstrate that compOG means cluster around 1 and CV is lower than that of non-compOGs. Non-compOG means cluster lower and CVs cluster higher.

To determine the impact of sparsely populated orthogroups, in addition to the selection of higher ploidy and parasitic taxa that may imbalance comparison between compOGs and non-compOGs, wilcoxon tests and Cliff’s Delta were used on subsets of data with and without orthogroups with mean protein numbers higher than 0.5 and parasites and higher ploidy taxa (Tab. 3). All tests supported the greater abundance of compOG proteins compared to non-compOGs, though exclusion of orthogroups with a mean number of proteins less than 0.5 significantly decreased this support.

**Table 3:**
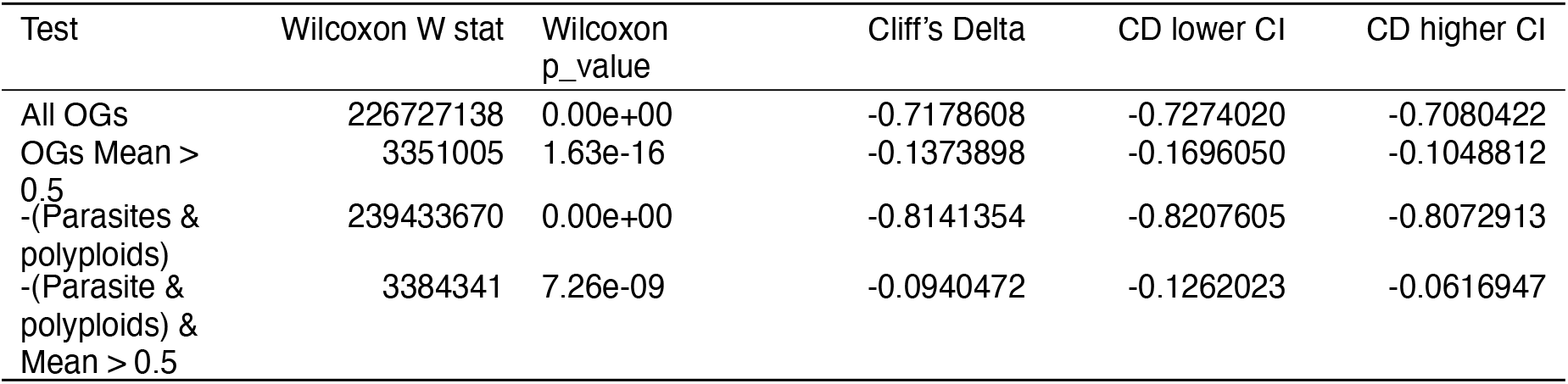
Support statistics for Cliff’s delta and Wilcoxon tests, comparing distributions of proteins across taxa for compOGs and non-compOGs. Wilcoxon p-values all support the alternative hypothesis that the mean number of proteins/taxa/compOG are generally shifted higher than the mean number of proteins/taxa/compOG. Cliff’s delta values were low, also supporting that compOG mean proteins/taxa/compOG were higher than OG mean proteins/taxa/compOG.

| Test | Wilcoxon W stat | Wilcoxon p_value | Cliff's Delta | CD lower CI | CD higher CI |
| --- | --- | --- | --- | --- | --- |
| All OGs | 226727138 | 0.00e+00 | -0.7178608 | -0.7274020 | -0.7080422 |
| OGs Mean > 0.5 | 3351005 | 1.63e-16 | -0.1373898 | -0.1696050 | -0.1048812 |
| -(Parasites & polyploids) | 239433670 | 0.00e+00 | -0.8141354 | -0.8207605 | -0.8072913 |
| -(Parasite & polyploids) & Mean > 0.5 | 3384341 | 7.26e-09 | -0.0940472 | -0.1262023 | -0.0616947 |

Proteins within compOGs tend to be more evenly distributed across taxa than other orthogroups, although this distribution may reflect genome and annotation differences. Only 28.1% of retrieved compOGs were among the most unevenly distributed orthogroups compared to 86.9% of non-compOGs (Tab. 4). Excluding the most unevenly distributed compOGs and non-compOGs, the remaining 18,182 non-compOGs were also less evenly distributed than the remaining 2721 compOGs (Fig. 2). CompOG means clustered near one, whereas non-compOG means clustered below one corresponding to considerable loss of non-compOG proteins in some taxa. Furthermore, compOG coefficents of variation are lower on average than non-compOGs, consistent with more even distribution; formal tests controlling for confounders are required to determine whether this pattern reflects dosage conservation.

**Table 4:** Unevenly distributed orthogroups. The vast majority of orthogroups retrieved in this study had proteins in few os the selected taxa, distorting the distribution plots. These orthogroups were ommitted from further analysis.

| Mean proteins/taxa/orthogroup | non_compOGs | compOGs |
| --- | --- | --- |
| 0.03225808 | 51592 | 493 |
| 0.06451613 | 63405 | 626 |
| 0.09677419 | 4144 | 108 |

To determine which compOGs and non-compOGs were universal, containing proteins for all queried species, orthogroups that did not map to any protein for any individual queried species were excluded. This resulted in retrieval of 86 universal non-compOGs and 184 universal compOGs, highlighting an overabundance of proteins that participate in the formation of protein complexes in the universal compOGs of this dataset. These results suggest that, although universal orthologs are rare across eukaryotes, they are likely to be dominated by compOGs.

Universal compOGs were manually assigned to functional groups linked to DNA, post-translational modification, RNA, translation and transport/structure mechanisms or other functions. For each compOG, I summarised per taxon counts in a heatmap with a cladogram of the taxonomic subset (Fig. 3). I then marked compOGs with counts at least 2.5 standard deviations above or below the across-species mean to identify expansions and reductions in 27 taxa. Expansion analyses excluded polyploid taxa (*P. tetraurelia, A. thaliana, A. macrogynus* and *E. huxleyi* ). Reduction analyses excluded parasites (*Encephalitozoon cuniculi, Plasmodiophora brassicae*, Apicomplexans). Notable expansions were identified in *Naegleria gruberi, Guillardia theta* and *Ectocarpus siliculosus*. No notable expanded compOGs were observed in parasites, consistent with known reduction and compaction of these genomes (Khalaf, Francis and Blaxter, 2024).

**Figure 3.**
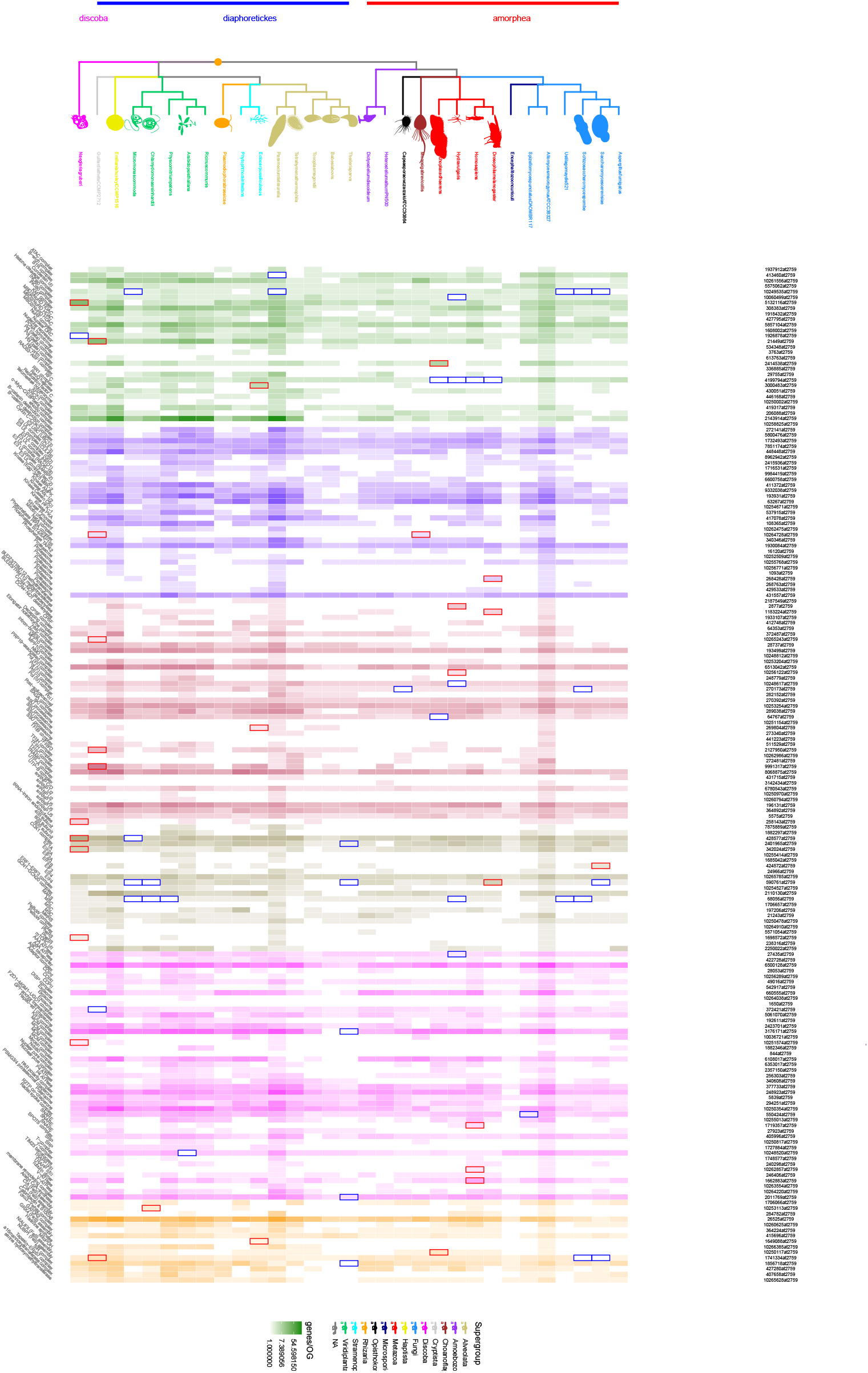
Repertoire of compOGs. Abundance of genes in 184 Universal CompOGs, stratified by functional category and by taxonomy. Intensity of squares corresponds to abundance of proteins per orthogroup for each species. Green squares correspond to orthogroups mapping to complexes with a role in DNA maintenance, purple corresponds to post-translational modifications (PTMs), maroon corresponds to RNA, beige translation, magenta transport/structure and orange for the remaining orthogroups. Expansions and reductions of orthologs within com-pOGs for individual taxa, relative to the other 30 taxa are represented as red and blue outlines, respectively.

Taken together, the results here suggest that the proteins of complexes present across eukaryotes can be binned into 184 groups by sequence. As proteins from each of these groups are present in every eukaryote queried, ancestral orthologs from each group can be inferred as present in the last eukaryotic common ancestor. Multiple taxon-specific expansions and reductions can be detected across the entire domain of eukaryotes. However expansions and stasis of other orthologs are observable across the domain and some individual supergroups.

### Quantifying the distribution of proteins across heterodimerically-related compOGs

Taxon-specific expansions and reductions of compOG proteins were apparent across the chosen eukaryotic subset. However, it was not clear how the number of proteins per compOG for taxa scaled in comparison to the number of putative corresponding interaction partners from other compOGs. To generate this data, I counted the number of proteins in each pair of compOGs that correspond to each physically determined heterodimer, inferring the extent of protein expansions and reductions of these

Heterodimer-forming proteins that each belong to different universal compOGs were found in 130 compOGs (Fig 4a). Summarised copy numbers of proteins in each universal compOG pair varied over 17-fold, from almost singlecopy (66) to 1170, suggesting broad heterodimer protein diversity across and within each species. Unexpectedly, no universal compOG pair contained exclusively single-copy proteins. However, most orthologs were not expanded more than two copies (median 145), suggesting evolutionary pressure to maintain fewer than three copies of heterodimer-forming proteins (bi-modal 72 and 126). Inferring that orthologs of heterodimer-forming proteins interact with corresponding orthologs, this data suggests that most eukaryotic heterodimeric interactions occur between one protein and two paralogous partners or between two pairs of paralogous partners (Fig 4c).

**Figure 4.**
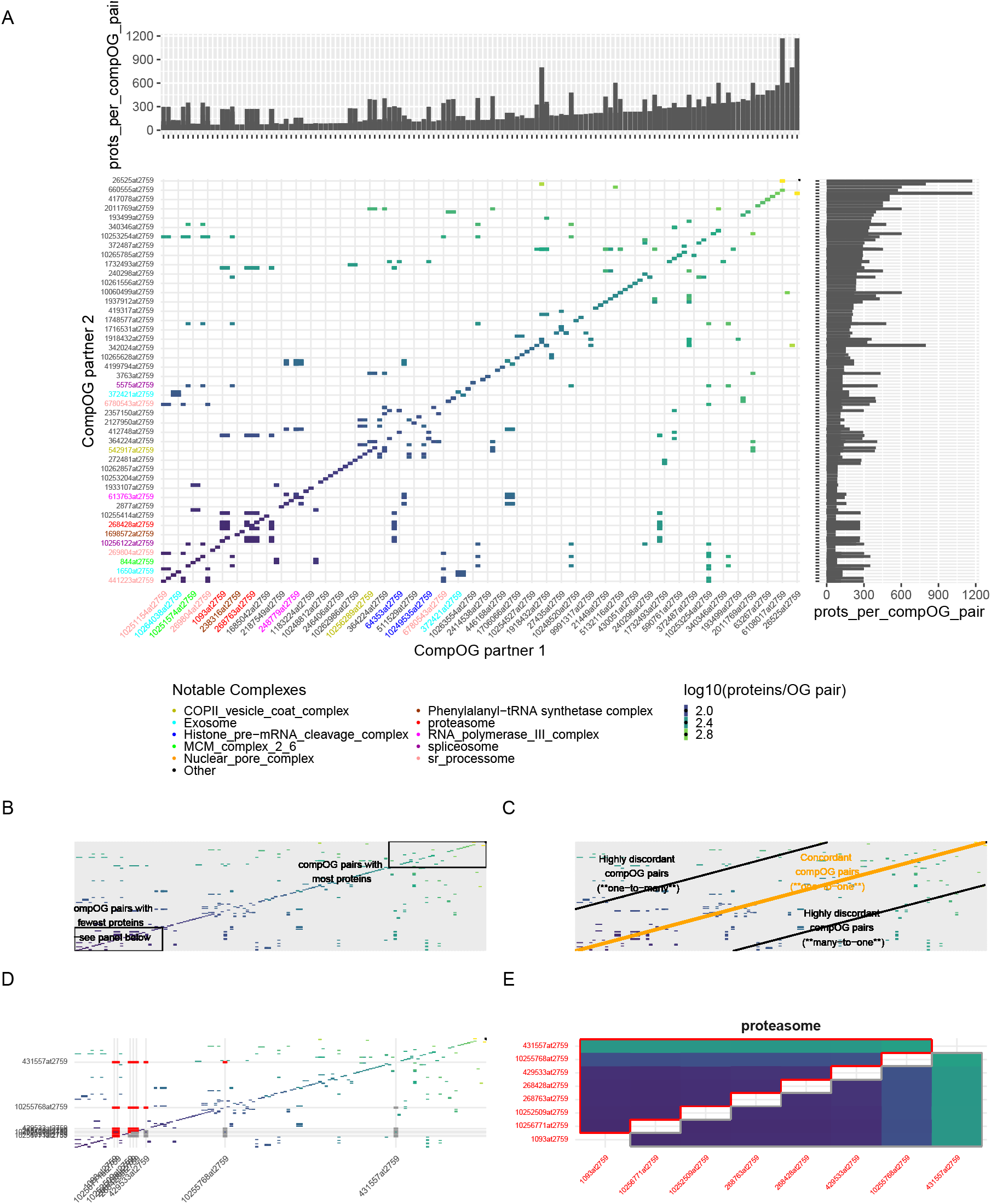
Repertoire of heterodimer-including compOG pairs, populated with abundance of proteins. Com-pOGs (axes labels) are ranked from fewest to most proteins from summarised protein counts for the 31 eukaryote subset. Each square intensity quantifies the number of proteins per compOG pair. Axes labels of every third compOG are shown for legibility (See Supplementary Figure S1 for all labels). Axes labels are shaded to reflect low-copy protein-containing macromolecular complexes to which compOG components belong. **A:** Reference map of pairwise counts of proteins in all-against-all universal compOGs.-The shape and color palette of this plot reflect a spectrum of duplicated orthologs, spanning multiple macromolecular complexes. Marginal histograms show number of proteins for partner 1 (top) and partner 2 (right). **B:** Low-copy compOGs (bottom-left box) predominantly contained components of macromolecular complexes but also occupied high-copy regions of the map (top-right box). **C:** Concordant pairs cluster near the diagonal, while discordant one-to-many and many-to-one pairs are offset from it. **D:** Proteasome pairs are highlighted to show the transition from concordant to discordant relationships. **E:** Proteasome-associated pairs are shown in isolation for direct comparison.

Despite this, many high-copy compOG pairs were also retrieved, interpreted as conservation of heterodimers that include one or two highly duplicated proteins across all eukaryotes. These compOGs included a compOG including Homeobox-like domains and Myb domains (2143914at2759) and an rbb4-containing histone-binding compOG (427795at2759) (Fig 4b). The single most expanded compOG pair was the myosin intermediate filament-containing compOG (6108017at2759) and the calmodulin A containing compOG, (26525at2759).

Many of the heterodimers retrieved in this study are components of macromolecular complexes. Categorising the heterodimers by the macromolecular complexes they participate in reveals that many macromolecular complexes contain interactions between single-copy proteins. The proteasome consists of 15 interactions from six (typically) single-copy proteins, presumed one-to-one interactions, an additional 13 interactions from two higher-copy compOGs and one presumed many-to-many interaction between the higher-copy compOGs (Fig 4d and e). This suggests that all such proteins can be determined from orthogroups and used as markers for new eukaryote genomes.

To investigate further, the least duplicated compOGs and all other compOGs belonging to the same macromolecular complexes were selected (Fig 5). As mentioned above, these compOGs corresponded to known components of the spliceosome, the proteasome, the ribosomal small subunit processome, MCM complex 2-6 and the exosome. The single lowest-copy compOG pair was 441223at2759 and 10251154at2759 corresponding to krr1 and WD-repeat containing protein 46 of the ribosomal small subunit processome, respectively.

**Figure 5.**
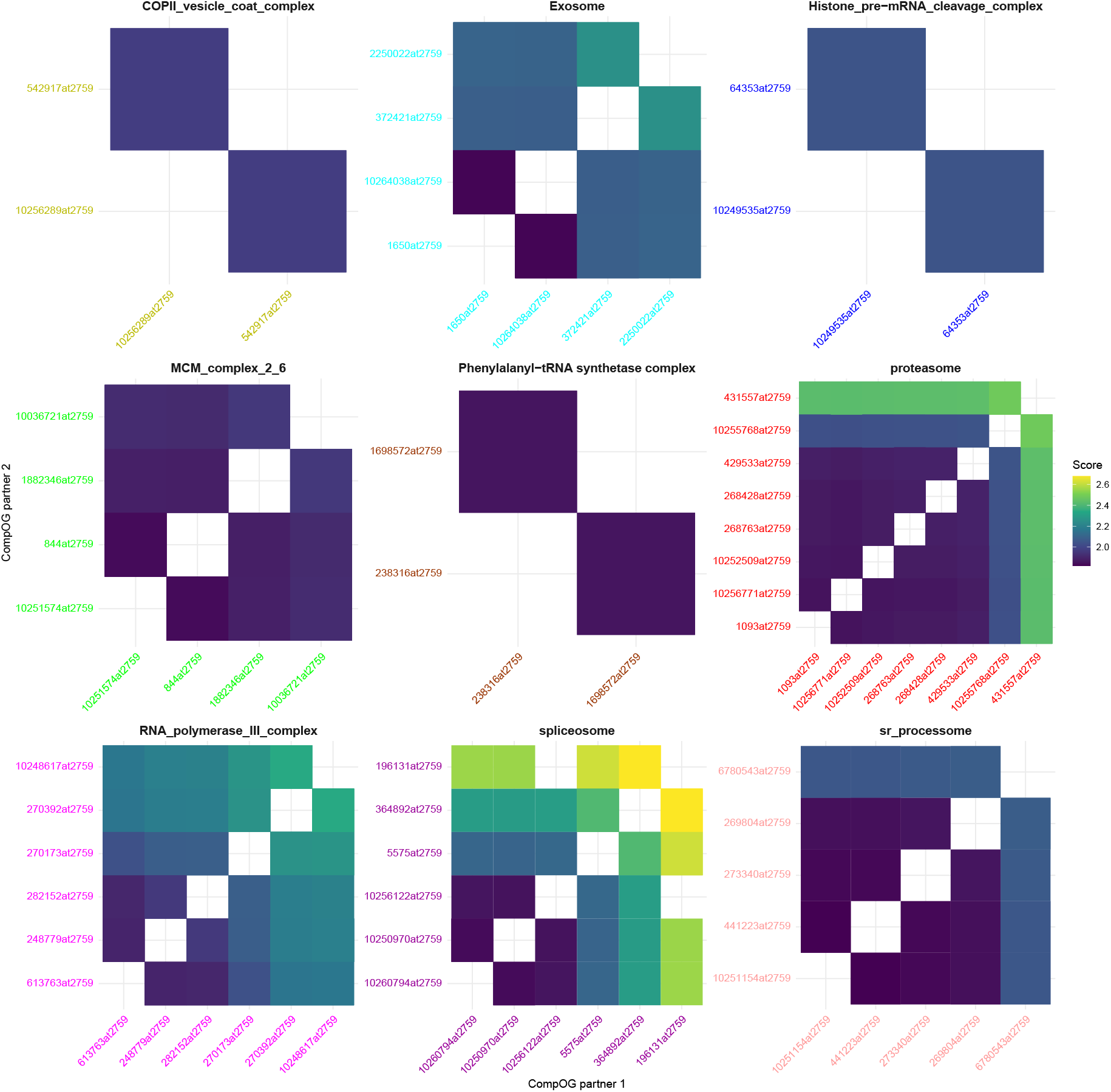
CompOGs corresponding to macromolecules that have the fewest proteins per species. CompOGs with the fewest proteins per species from the heterodimeric interaction heatmap, stratified by macromolecular complexes they belong to. The same colour scheme is used.

Notably, the compOGs corresponding to subunit 2 of splicing factors 3A and 3B, of the spliceosome, were the least expanded, universally-represented compOG pair. The phenylalanine–tRNA ligase, comprised of subunits alpha and beta compOGs, represent the complex with the least expanded OGs.

These results reveal that at least nine macromolecular protein complexes are conserved in all queried eukaryotes, the last eukaryotic common ancestor and that multiple heterodimers that participate within these complexes are resistant to duplication events. Identification of the low copy components of these complexes alone rationalised identification of corresponding higher copy components. The presence of multiple low-copy components in many of these macromolecular complexes suggest that single-copy components of macromolecular complexes may be used as completeness markers for sequencing of new eukaryotic genomes.

### The expanded complex-forming orthogroups of *Naegleria gruberi* and *Guillardia theta*

The number of proteins in each universal compOGs, that is the total number of universal, protein-binding orthologs summarised over the selected eukaryotic taxa, is a useful generalisation of the number of copies of corresponding complex-forming proteins. However, complex-forming proteins encoded in each individual eukaryotic genome are likely to vary in scale from this consensus, which may indicate divergent mechanisms within specific taxa. I therefore sought to count the number of proteins per compOGs of individual taxa, reusing the ranks of protein counts per compOGs to determine taxon specific deviations of protein-binding orthologs from all other queried eukaryotes.

Protein complexes of *Naegleria gruberi* and *Guillardia theta* are of key interest. Phylogenomic evidence frequently places excavates, like Naegleria, near the root of the eukaryotic tree of life, strengthening the hypothesis that the last eukaryotic common ancestor was an excavated eukaryote (Jewari and Baldauf, 2023; Williamson *et al*., 2024). *G. theta* is an algal cryptophyte that, like other cryptophytes, harbours four simultaneous genomes from four compartments; a nucleus, a nucleomorph, a plastid and a mitochondrion (Curtis *et al*., 2012). Analysis of the nuclear genome of *G. theta* has revealed endosymbiont-nuclear transfer.

With a clear signal that compOG expansions have occurred in both lineages (Fig 4), I sought to investigate this further, comparing pairwise protein numbers of heterodimeric-binding partner containing compOGs. For *N. gruberi* and *G. theta*, rank-order of compOGs in x and y-axes of the heatmap, therefore the location of co-ordinates, again described the spectrum of expected abundance of proteins (based on the sum of numbers of proteins per compOG for the subset of 31 eukaryotes). However, the colour of co-ordinates and the size of the marginal histogram bars describe the spectrum of observed number of proteins per compOG pair (Fig 6a and b, respectively). This time I marked compOG pairs that were more than a combined standard deviation (sd for compOG partner 1 + sd for compOG partner 2) away from the combined mean for compOG pairs.

**Figure 6.**
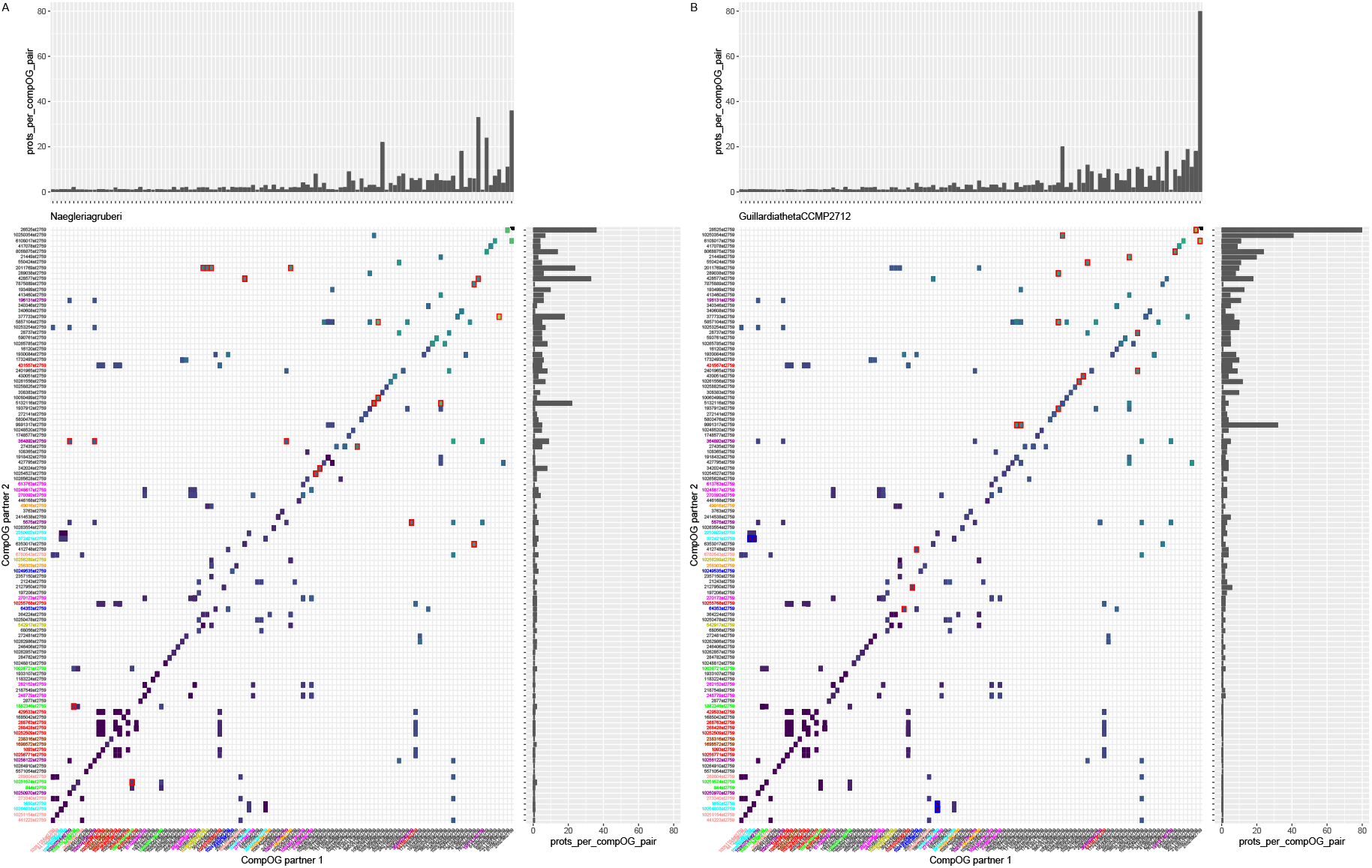
Comparison between pairs of universal compOGs that correspond to heterodimers from pan-eukaryotes and either *Naegleria gruberi* or *Guillardia theta*: **A. Comparison between universal compOG pairs of paneukaryotes and *N. gruberi*.** CompOG heatmap order is retained from Fig. 4 and co-ordinate positions are defined by the number of proteins per compOG for all 31 eukaryotes. Co-ordinate colour is defined by the number of proteins per compOG for *N. gruberi*, also displayed as margin histograms in the top and right-hand margins. Expansions and reductions of protein numbers per orthogroup pair in *Naegleria* are identifiable by cross-referencing bars in the marginal histograms, co-ordinate colour and co-ordinate location. Heatmap co-ordinates corresponding to large histogram bars in the top and right-hand margins, that have yellower colours (2011769at2759, 428577at2759, 7875889at2759, 10060499at2759, 5132116at2759) or are flanked by bluer colours (342024at2759, 1706657at2759, 10251574at2759) are expansions, outlined in red. Co-ordinates that correspond to reductions are identifiable by a darker colour, flanked by yellower colours (1918432at2759, 2250022at2759) and are outlined in blue. **B. Comparison of heterodimeric interactions between universal compOGs of *Guillardia* and pan-eukaryotes**. Inverse compOG pairs are not outlined in this figure.

For *N. gruberi*, the pairs of compOGs that were highlighted as containing more proteins compared to other taxa were driven by expansions in three high-copy compOGs, 428577at2759 (type-i, ubiquitin-like domain-containing protein), 5132116at2759 (actin) and 2011769at2759 (ADP-ribosylation factor), 342024at2759 (elongation factor 1-alpha) and 364892at2759 (snRNP U5) containing compOGs, two duplicated commonly single-copy compOGs, 10251574at2759 (DNA replication licensing factor) and 1698572at2759 (phenylalanine–tRNA ligase subunit beta). Conversely, the single *N. gruberi* protein in the 1918432at2759 (MBD2/NuRD HDAC) compOG drove the single notable reduction, specific to *N. gruberi*. These expansions suggest *N. gruberi* specific expansions of putative components of the MCM complex, the phenylalanine tRNA synthetase and spliceosome, macromolecular complexes that are central to three intricate cellular mechanisms.

For *G. theta*, four high-copy compOG expansions contributed to highlighted outlier compOG pairs. Two high-copy compOGs, 26525at2759 (calmodulin-like protein) and 9991317at2759 (TFIIIC-like transcription factor), corresponded to non-significantly expanded compOGs. The remaining high-copy compOG expansions, 8068875at2759 (TRAMP complex protein) and 10261556at2759 (BTR complex protein), each corresponded to an additional expanded compOG, 64767at2759 (SKI complex) and 430051at2759 (RecQ helicase), respectively. Lower-copy compOG expansions included 2127950at2759 (TFIIB/D-type transcription factor), 2187549at2759 (Trm112 methyl-transferase) and 412748at2759 (Cleavage and Polyadenylation Specificity Factor). The RNA exosome component compOG, 372421at2759 (DIS3 nuclease) was the single-copy compOG that contributed to the reduction of proteins in three compOG pairs.

In summary, these results pinpoint the taxonomic expansions and reductions of specific groups of heterodimers from the genomes of multiple eukaryotes. Retrieval of taxon-specific expansions and reductions from existing orthology data was a necessary step for selection of expansion-rich taxa. However, pairwise predictions of ortholog binding networks revealed expansions within the INO80 complex, the ribosome and the membrane modeling complex of *N. gruberi* and considerable expansion of the calmodulin-like, EF-hand proteins in the complexes of *G. theta*. That these species were identified from such a diverse subset, reinforces the potential of using orthogroups, specifically compOGs, to investigate discordant protein expansions and reductions across eukaryotes.

## Discussion

The results of this study suggest that protein complex components and their orthologs are less susceptible to loss of corresponding genes than all other eukaryotic orthologs, which justifies comparison of numbers of proteins across orthogroups and taxa. Although only 184/82,213 orthogroups included proteins from each query eukaryote and any protein complex component, those universal orthogroups are sufficient for discerning taxon-specific and taxon-independent differences in the number of proteins per orthogroup. Presence of compOG proteins across all 31 species spanning 11 supergroups support broad conservation (Fig. 3,4 and 5). Furthermore, 130 of these compOGs include proteins that form complexes with proteins from other orthogroups of this repertoire. Counting and plotting the number of proteins in each pair of these orthogroups, allows characterisation of the number of potential interactions between copies of protein-binding orthologs and detection of species-specific duplications or losses of these proteins.

Characterisation of proteins across the taxa of this study suggested that most universal compOGs interact with at least one compOG containing a similar number of proteins and a compOG containing a different number of proteins. The resulting distribution can be inferred to represent heterodimeric pairs applicable to all eukaryotes and can be used as a framework of expected proteins per compOG pair (in space) with which to compare observed proteins per compOG pair for an individual species (colour). Applying the spatial repertoire of 130 compOG pairs to *Naegleria gruberi* and *Guillardia theta*, species-specific expansions were identified in the context of binding partners.

The even distribution of proteins within compOGs across taxa and the higher number of compOGs containing proteins from all taxon queries (68%), both compared to non-compOGs, suggest that homologs of eukaryotic protein complex components are ancestral proteins that are generally less prone to taxon-specific loss than other proteins. These results may be a result of variation in genome completeness, ploidy, and annotation quality. Although known higher ploidy taxa and parasites were explicitly excluded from taxon-specific quantification of protein duplications and reductions respectively, normalised dispersion metrics were not estimated for these covariates in this study.

Retention of protein complex component homologs could be explained by underlying pressure within extant and ancestral eukaryote genomes to maintain protein-level dosage of protein complexes by fixing the genes coding for protein complex components and regulatory elements required for their expression. Papp et al hypothesised that an imbalance (disproportional expansion or reduction) of heteromeric protein complex components are likely to be deleterious, supporting the hypothesis with the observation that most dosage-sensitive genes identified from *Saccharomyces cerevisiae*, code for protein complex components (Papp, Pál and Hurst, 2003). Further experiments in *S. cerevisiae* demonstrate fitness costs attributable to duplication of different heteromers, though such observations may be specific to the mutational burden of the specific model (Diss *et al*., 2014, 2017; Veitia, 2017).

Duplications of putative protein complex component paralogs can be inferred from the presence of multiple proteins from each taxa within individual universal compOGs. Although these proteins may not necessarily act within the same protein complexes, paralogs of protein complex components may contribute to cellular robustness and buffer functional instability, participating as replacement components when cognate components are mutated, insufficient or absent (Dandage and Landry, 2019). Heterodimeric relationships between compOGs with many and few proteins (many-to-one and one-to-many, Fig. 5) or between compOGs with even (one-to-one and many-to-many) may correspond to dosage-insensitive and dosage-sensitive genes, respectively. Although both categories of orthogroup are clarified using the strategy outlined in this study, future analysis should implement a stoichiometry metric to predict comparative buffering capacity of each proteome for tolerating deletion of proteins from the compOG pair repertoire. It is important to acknowledge that post-translational mechanisms of stoichiometric control also contribute to protein-level dosage compensation (Ishikawa, Ishihara and Moriya, 2020).

Taxon-specific protein losses were inferred from the majority of orthogroups, reflecting the tendency of genes to be lost throughout evolution (Lynch, 2007). For emergence of protein-binding paralogs, the evolutionary pressure to fix single-gene duplications rarely overcome the power of drift, in part because of the requirement for protein-level dosage compensation and many duplicates never reach fixation (Lynch, 2007). Although this transience is partially overcome by dosage-preserving, whole-genome duplications, this may suggest that proteins from universal orthogroups co-evolve together and could serve a role as phylogenetic markers.

To infer the extent of protein complex component duplication across eukaryotes, the PCOC suite of scripts was created from the repertoire of universal compOGs. PCOC generates maps corresponding to the number of proteins within universal compOGs, stratified either by the taxonomic subset used in this query or by compOGs containing heterodimeric binding partners of proteins within each compOG. General patterns of protein distribution within the maps suggest some properties of compOG proteins.

First, the compOGs with the most proteins, inferred as the most duplicated complex component orthologs, are readily identifiable. These include components of the INO80 complex (10060499at2759), R2TP co-chaperone (10250354at2759), MBD2/NuRD HDAC (427795at2759), Calprotectin & S100A8 (26525at2759) and Myosin complex (6108017at2759). These compOGs have main functions in post-translational modifications, transcription factors, transport/structure and, in the case of EF-hand proteins, calcium sensing perhaps suggesting that some buffering of dosage sensitivity may exist within these mechanisms in eukaryotes.

Ancestries of EF-hand proteins (like calmodulin A) have been widely reported as extensive, multi-copy in LECA and present in ancestral and extant bacteria (Nakayama and Kretsinger, 1994). The expansion of such proteins may reflect adaptive pressure incurred by an increased repertoire of targets. Folding of an increased repertoire of proteins, binding an increased repertoire of DNA enhancers/promoters and sequestering calcium for an increased signaling network are all feasible pressures that may have driven the expansion of these proteins. Exceptions were clearly outlined in the maps for further analysis. One example was the compOG with the most proteins of the repertoire, the calcium-calmodulin-myosin complex compOG (26525at2759), that included multiple proteins for all species and up to 105 copies in Emiliania huxleyi, but only two copies in Saccharomyces cerevisiae.

Second, universal compOGs with the fewest proteins were readily identifiable and corresponded to the least duplicated families. These included components of the SSU processome (10251154at2759 and 10251154at2759), exosome (10264038at2759 and 1650at2759) and snRNP U2 (10250970at2759) which are associated with processing of the small subunit of ribosomes (SSU), transport and spliceosome. No taxon had a statistically-significant expansion or reduction of these compOGs. Although some higher-copy ribosome, exosome and spliceosome compOGs were identified in this study, the overall pattern suggests that ribosome, exosome and spliceosome machinery are sensitive to protein-level dosage. As an example, dosage imbalance of ribosomal components may reflect a risk of impeded protein folding. Proteinopathies, typically characterised by aggregations of misfolded proteins, are well known in the context of human and animal disease and ageing. The impact of the ribosome has been hypothesized in these and other conditions (Polymenis, 2020; Stein *et al*., 2022; Iben, 2023).

Most universal compOGs had low coefficients of variation (100/184 < 0.6) and means near one, which further suggest resistance to duplication. Accordingly, 53/184 universal compOGs (20/86 universal non-compOGs) had a CV < 0.6 and a mean between 0.95 and 1.5. As has been demonstrated by this study, low-copy compOGs may comprise an additional metric for assessment of eukaryotic genome, transcriptome and proteome completeness. Only 9 of the 44 compOGs populated with the fewest genes are also in the eukaryota Benchmarking Universal Single-Copy Ortholog (BUSCO) dataset. This reflects comparatively restrictive conditions of the BUSCO dataset, but does not negate the usefulness of compOGs, BUSCOs or both metrics for sequence analysis.

Third, maps highlighted compOGs that had a number of proteins two standard deviations greater than the mean, inferred as taxon-specific duplications of protein complex components. These expansions were observed in *Naegleria gruberi, Guillardia theta, Ectocarpus siliculosus, Chlamydomonas reinhardtii, Monosiga brevicollis, Trichoplax adhaerens, Hydra vulgaris, Homo sapiens*, Drosophila melanogaster and Saccharomyces cerevisiae. Although a significant number of expansions corresponded to Metazoa, most taxon-specific expansions occurred in the diaphoretickes, excluding the WGD-driven expansions of viridiplantae and Emiliania huxleyi.

These results suggest an increase in paralog number in the last common ancestor of diaphoretickes (viridiplantae, haptophyta, cryptophyta and the SAR group), that may be a result of ancestral whole genome duplication. However, reference organisms, plants and ciliates are not necessarily representative of diaphoretickes. Further analysis was limited to *N. gruberi* and *G. theta* compOG expansions, although future analyses of the remaining taxon-specific expansions highlighted here may provide insight into further duplications of eukaryote protein complex components.

Naegleria gruberi expansions were identified for actin-related proteins, CLP1-related proteins, ribosomal proteins, eukaryote translation initiation factor proteins, phenylalanine-synthetases and MCM proteins. Interestingly, this includes expansions of two expected single-copy protein-containing compOGs, hypothesised as putative keystone components of macromolecular complexes.

Aminoacyl-tRNA synthetases (ARSs), like Phenylalanine-tRNA synthetase (PheRS), are essential proteins that conjugate amino acids to their respective transfer RNAs. They are ubiquitous across eukaryotes and have been used in early eukaryotic tree of life estimations (Brown and Doolittle, 1995). Although variants of ARSs are found in eukaryotes, archaea, bacteria and some viruses, ARS-containing compOGs only contained eukaryotic ARSs that map to the host nuclear genome. Eukaryotic ARSs impact their hosts as part of multisynthetase complexes (MSCs), which can include paralogs, such as the MSC-bound, long form of arginyl-tRNA synthetase in mammals which exists alongside an N-terminal truncated shorter form (Yang *et al*., 2014).

N. gruberi has two Phenylalanine-tRNA synthetase (PheRS) beta subunit genes, unlike most eukaryotes queried in this study, which have one. There is a clear size difference between each of the pairs of PheRS proteins, suggesting the existence of free and MSC-bound forms. However investigation of the expressed forms of these genes has not been reported. Furthermore, although not included in this study, the pathogenic relative species, Naegleria fowleri, also harbours multiple isoleucine tRNA synthetases. ARSs have been proposed as therapeutic targets and any subfunctionalisation through multiple ARSs should be fully investigated (Tillery *et al*., 2021).

The minichromosome maintenance (MCM) complex is a DNA helicase, integral for DNA replication. MCM proteins first emerged as a single protein, comprising a homohexameric complex in an Archaeal ancestor of eukaryotes (Liu, Richards and Aves, 2009; Chia, Cann and Olsen, 2010). MCM family proteins are conserved across eukaryotes, therefore inferred to be conserved in LECA. The evolutionary distance between archaeal MCM and eukaryote MCM protein families suggest a considerable amount of undocumentable change between LECA and the first eukaryotic common ancestor (FECA), that may include the diversification of the MCM proteins.

Naegleria compOGs included more MCM proteins, particularly MCM3 and 4 than other eukaryotes. Previous analyses by Liu et al found duplicate MCM3 in Naegleria and no other protists via BLAST searches, but no duplicate MCM4 was identified in this study in Naegleria or any other species. MCM3 is also duplicated in Xenopus laevis and Danio rerio, hypothesized to have been duplicated in the ancestor of all vertebrates and subsequently lost from descendants (Shinya *et al*., 2014). Duplicate MCM3 and 4 proteins in Naegleria suggest that LECA had an expanded repertoire of MCM proteins. Loss of MCM proteins may have contributed to diversification of early eukaryotes. Incorporation of orthology and protein interaction data can therefore signpost proteins such as these for future phylogenetic analysis.

Expansion of actin-related proteins has been acknowledged in Naegleria (Dawson and Paredez, 2013). Although LECA could have had an expanded actin-related protein repertoire, like Naegleria, another possibility is that expansion of ARPs occurred specifically within Naegleria. Further high-quality, free-living discoban genomes are required to investigate further.

CompOG expansions in the Guillardia theta nuclear genome were also of interest because of the presence of nucleomorph, mitochondrial and plastid genomes in the cryptophyte. G.theta has expansions in transcription factors, TRAMP, SKI and BTR complex components and EF-hand proteins. No expansion of the expected single-copy macromolecular complex components was observed here, although one expected single-copy compOG expansion that was unassigned to a macromolecular complex was observed (2187549at2759).

Notably, *G. theta* only has one copy of DIS3, the catalytic nuclease component of the RNA exosome. Deviations from canonical exosome repertoires are known, but complete absence of DIS3 paralogs is uncommon. In this dataset, only the heavily reduced and compacted microsporidian *E. cuniculi* has a single DIS3 protein. Divergent repertoires have been reported for *Plasmodium falciparum* and *Trypanosoma brucei* (Droll *et al*., 2018; Cesaro *et al*., 2023), but neither species are restricted to one protein in this orthogroup (data not shown - UniProt search for taxa and 372421at2759). As some functional redundancy has been reported in human DIS3, DIS3L and Rpr44, it follows that the single DIS3 homolog in *G. theta* may undertake the molecular roles of each of the DIS3 paralogs in other eukaryotes. Nevertheless, these findings outline these protein families as an area of further study in *N. gruberi* and *G. theta*.

There are a number of clear limitations to the approach of domain-wide investigations of protein complexes at orthogroup level, in the context of both the orthology data and the protein interaction data used.

One limitation of the orthology data used in this study is the taxonomic breadth of species available from the database. The version of OrthoDB used (odb12v0) includes 5,827 eukaryote species, outnumbering the numbers of eukaryotes in the OMA (789) and the evolutionary genealogy of genes: Non-supervised Orthologous Groups (EggNOG - 1322) databases (Hernández-Plaza *et al*., 2022; Altenhoff *et al*., 2024; Tegenfeldt *et al*., 2025). However, there are many other unrepresented eukaryote supergroups absent from this database largely because of the absence of corresponding genomes (Schoenle *et al*., 2025). Inclusion of additional protist species, particularly heterotrophs that have been recently sequenced, was not possible for this study. Further investigation of the Complex Portal across eukaryotes would optimally include published Hemimastigophorean, Provorean, Metamonada and Malawimonada genomes.

Another limitation of the orthology data is the inability to resolve some paralogs. Generally, orthogroup level analyses are incapable of resolving isoform or paralog specific conservation patterns for orthogroups containing multiple paralogs. Protein-level resolution and phylogenetic methods beyond those currently used to create orthology databases are required to discern full hierarchies of orthogroups or determine which proteins have undergone subfunctionalisation or neofunctionalisation relative to an ancestral paralog (Ohno, 1970).

A limitation of the protein data used in this study is the narrow taxonomic representation of the queries that comprise the database. The EBI Complex Portal is largely limited to physical protein interaction data from experiments using a narrow selection of reference eukaryotes; *Homo sapiens, Mus musculus, Saccharomyces cerevisiae, Drosophila melanogaster, Arabidopsis thaliana, Caenorhabditis elegans, Schizosaccharomyces pombe, Rattus norvegicus* and *Gallus gallus*. These species are among the most established eukaryotic references, each corresponding to accessible genomes, gene predictions and research material such as antibodies and cell lines facilitating these findings. Future implementations of the approach used in this study will benefit from any additional protein complexes curated in the EBI Complex Portal as additional eukaryotic models become amenable to protein complex identification. Alternatively, lowering the threshold of evidence required, such as including co-association data, is another mitigation strategy to the issue of narrow taxonomic breadth of input data.

There are also limitations of the scope of this study. First, inclusion of parasites in this subset is likely to restrict the number of retrieved universal orthogroups below the actual repertoire of LECA. However, this decision filters proteins amenable to loss, increasing the predictive power and overall likelihood that proteins corresponding to the repertoire will be encoded in any eukaryotic genome of interest. Therefore, it is possible that a different number of universal compOGs are predicted from orthology mapping the same protein interaction data used in this study to different combinations of eukaryotes. It is possible that many of the same orthogroups from the repertoire predicted in this study could be retrieved from different taxa that span the same eukaryotic supergroups. Future analysis will aim to confirm this.

Second, the use of any orthology database mapping restricts analysis to the comprehensiveness of the tool used to create the database. The presence of trans-splicing mechanisms in some lineages (euglenozoa for example) can hinder protein prediction (Günzl, 2010; Kostygov *et al*., 2024). For protein complex predictions in taxa with non-canonical splice mechanisms, fully comprehensive, sequence-homology search methods should be used, either in the orthology database construction or instead of an orthology mapping approach. However, use of orthology and protein-interaction data is a faster, more accessible approach that can augment formal ancestral reconstruction.

Third, this study does not map orthology to prokaryotic organisms. The mosaic nature of LECA, an organism consisting of multiple prokaryotic ancestors justify inclusion of prokaryotic genomes in addition to extant eukaryotic genomes for the broadest possible comparative analyses, though that falls outside of the scope of this study. Future homology searches using the protein Complex Portal should query archaeal genomes from the Thaumarchaeota, Aigarchaeota, Crenarchaeota, and Korarchaeota (TACK) group, considered antecedent to LECA. It may be the case that the Complex Portal includes archeal protein complex data in the future. Indeed, the Complex Portal includes some bacterial and SARS-CoV-2 protein complexes, though they were not used as queries in this study.

Technical considerations include the use of reference genomes, rather than transcriptomes, to predict a repertoire of protein complexes. This decision ensures that results can be used for downstream comparative analysis of eukaryote genomes, such that presence or absence of proteins in the repertoire is not impacted by expression. However, the repertoire may be used to validate completeness of transcriptomes. A vast array of transcriptomes are now available and the additional search space offered by transcriptomes, particularly of under-represented protists, is of interest for deconvolving relationships within newly described clades and across the entire domain of eukaryotes.

Finally, species-specific variation may also affect how accurately these results reflect the true eukaryote pan-complexome. It is possible that false negatives, unobserved protein complexes, or false positives, such as species-restricted complexes exist in the proteomes of this dataset, as well as species-restricted complexes occur in organisms that have not yet undergone proteomic analyses.

In summary, the results here provide the research community with a list of 184 orthogroups for broad comparative analysis of protein complexes within and across eukaryotes. Although formal ancestral reconstruction is required for confirmation, it is highly likely that proteins corresponding to all 184 compOGs were present in LECA and supported by the observation that the universal compOGs with the fewest proteins contain proteins from macromolecular complexes widely predicted to be in LECA as well as nearly all extant eukaryotes. The dominance of compOGs over other universal orthogroups suggest a necessity of protein complex components over other proteins and similar numbers of proteins within heterodimerically-related compOGs reinforce the importance of protein-level dosage compensation. The publication of this repertoire of compOGs complements the available set of tools for comparative analysis of ever-increasing eukaryotic sequence data.

## Acknowledgements

This work was funded by a Wellcome Sanger Excellence Fellowship. I would like to thank Eduard Ocaña-Pallarès, Mark Blaxter, Michael Ansah, Charlotte Wright, Merce Montoliu Nerin, Lewis Stevens, Marcela Uliano-Silva, Jessica Thomas, Richard Challis, Claudia Weber, Amjad Khalaf, Cibele Sotero-Caio, Erna King, Martha Mulongo, Witold Morek, Yan Liang, Maggie Georgieva, Julien Martinez, Hend Abu-Elmakarem, Richard Durbin, Sucharitha Balu and Sandra Orchard for their helpful comments on the manuscript.

## Supplementary Data

**Figure S1:**
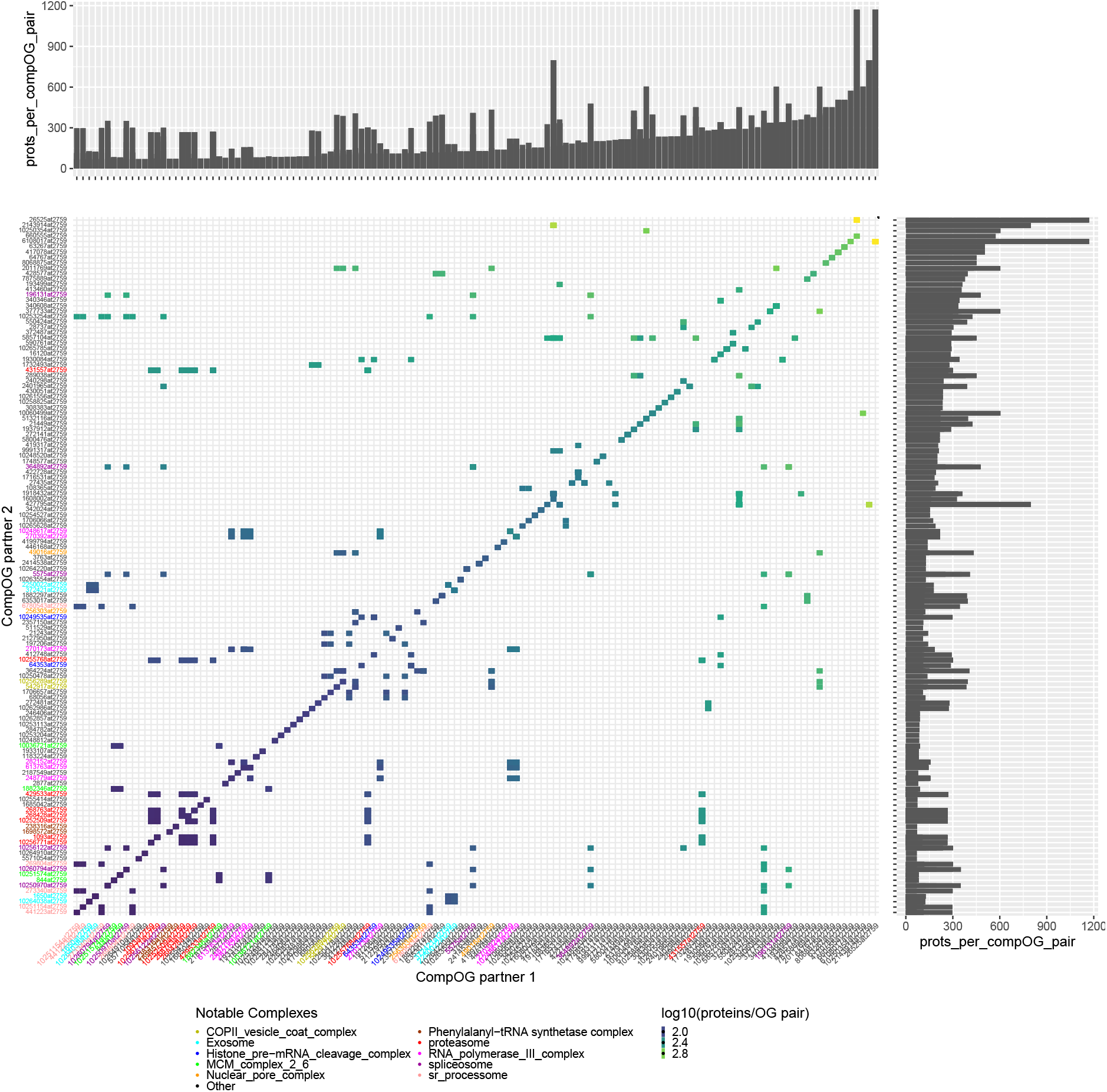
Supplementary Figure 1. Full compOG pair matrix

